# Oxalotrophic bacteria in desert and drylands: Enzymatic pathways and Carbon sequestration

**DOI:** 10.1101/2025.02.10.637404

**Authors:** Arun N. Prasanna, Sabiha Parween, Katja Froehlich, Tanja Schmidt, Kirti Shekhawat, Durga D. Prabhu, Maged M. Saad, Heribert Hirt

## Abstract

Oxalotrophy refers to the ability of bacteria to utilize oxalate as a carbon and energy source. This is a critical process with significant implications for the global carbon cycle. Oxalate-degrading bacteria play a key role in carbon sequestration through the oxalate-carbonate pathway (OCP), contributing to stable inorganic carbon pools. In this study, we identified and cataloged 20 enzymes associated with various facets of oxalate metabolism to characterize the oxalotrophic potential of bacteria. Within this group, sets of enzymes were grouped into two functional categories in the context of carbon sequestration: a biomineralization toolkit for converting oxalate to inorganic carbon and an assimilation toolkit for incorporating oxalate into metabolic pathways such as amino acid biosynthesis and energy production. Using bioinformatic approaches, we analyzed a collection of 536 bacterial genomes from desert and dryland strains spanning 81 genera to identify oxalotrophs. To validate our findings, we tested several bacterial strains for growth on media supplemented with exogenous oxalate. Notably, while multiple bacterial strains grew on oxalate media, two Pseudomonas species, namely JZ043 and JZ097, failed to grow despite genomic predictions suggesting otherwise. Further investigation of these strains revealed several non-conservative amino acid substitutions in the glyoxylate carboligase enzyme (EC 4.1.1.47), a key player in oxalate metabolism, suggesting a potential link between these mutations and their inability to metabolize oxalate. Our findings highlight the significance of our approach for identifying oxalotrophic bacteria (OxB) and offer valuable insights into the molecular basis of oxalate metabolism.

**Graphical Abstract:** 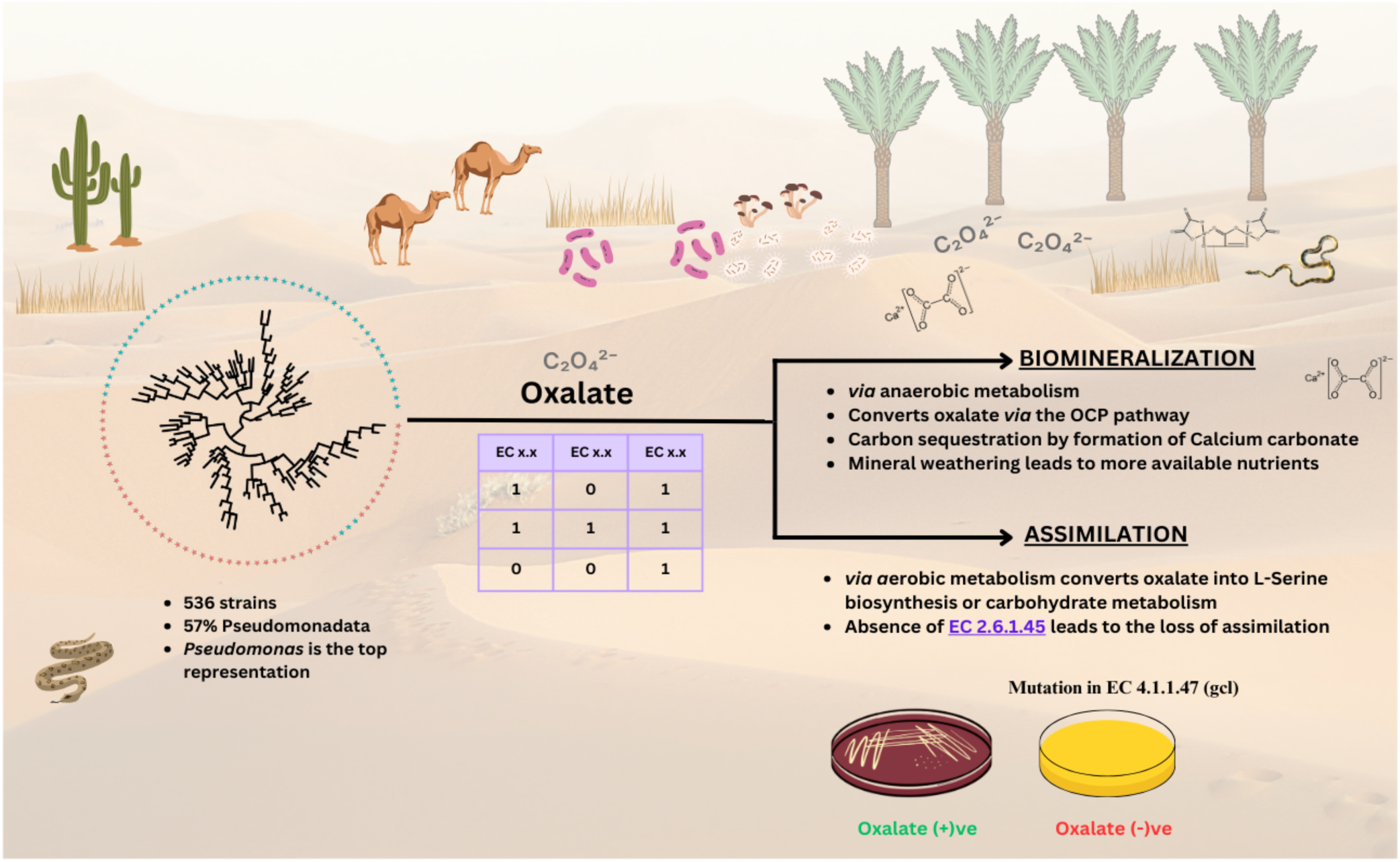

## Introduction

Arid soils are formed by extreme dry climates and typically have a sandy, coarse texture, resulting in low water retention and low organic matter content. This limits their soil fertility and reduces their ability to support vegetation (Naorem et al. 2023). Besides, arid soils are often highly calcareous and contain abundant calcium (Ca²⁺), challenging plants to adapt for survival. Calcicole (chalk-loving) species excel in such environments by tolerating elevated calcium levels and accessing nutrients with low solubility (Lux, Kohanová, and White 2021). To survive under these conditions, plants use key strategies including restricting calcium uptake through apoplastic barriers or sequestering excess calcium as calcium oxalate crystals in specific cell types. This sequestration prevents calcium toxicity and maintains ionic balance, highlighting calcium’s vital role in plant adaptation to calcareous soils (Nakata 2003). Despite employing these strategies, plants are not always successful, making the challenges of high calcium levels a critical concern for agriculture and future food production (Hirt et al. 2023). To address these challenges, we explore the role of oxalate-degrading bacteria (OxB) in sustainable agriculture and enhancing soil carbon sequestration.

Oxalic acid is a ‘Swiss army knife’ among the existing low molecular weight organic acids produced by plants, fungi, animals, and bacteria. It plays an important role in various environmental processes, such as soil weathering (Aragno and Verrecchia 2012; Schmalenberger et al. 2015), nutrient cycling (Pons et al. 2018), carbon sequestration through the oxalate-carbonate pathway (OCP) (Aragno and Verrecchia 2012), promoting plant growth (Kumar, Irfan, and Datta 2019), environmental detoxification with metal chelation, bacteria-fungi interaction (Palmieri et al. 2019), and land reclamation with application spanning environmental sustainability to human health (Duncan et al. 2002; Daniel et al. 2021; Ermer et al. 2023). Oxalic acid, a key component of root exudates, recruits oxalotrophic bacteria, which play an essential role in carbon cycling and plant growth promotion.

Oxalotrophy has been recognized as an ecosystem service owing to its contribution to carbon cycling (Cowan et al. 2024). Since the isolation of the first oxalate-degrading bacterium from the rumen of sheep grazing on oxalate-rich plants (Allison et al. 1985), significant advances have highlighted their ecological and physiological importance. For instance, in *Burkholderia* species, oxalotrophy is exclusively linked to plant growth-promoting strains and is essential for successful plant colonization (Kost et al. 2014). Fungal pathogens such as *Botrytis cinearea* secrete oxalic acid as a pathogenicity factor to infect their host. *Cupriavidus campinensis* degrades oxalate rendering protection from infection by up to 70% in Arabidopsis, tomato, cucumber, and grapevine (Schoonbeek et al. 2007). Furthermore, oxalotrophic bacteria (OxB) have been proposed as probiotics for treating hyperoxaluria and kidney stones in humans (Karamad et al. 2022).

Oxalotrophy is not a universal trait in bacteria. The capability depends on the presence of specific metabolic pathways and enzymes. Two models have been proposed for oxalate metabolism: one involves the conversion of oxalate to CO_2_ *via* anaerobic metabolism (Daniel et al. 2021). The generated CO_2_ either diffuses passively or actively through an active transporter out of the cell to precipitate calcium into calcium carbonate (Xia et al. 2024). Key players in this OCP pathway are the enzymes formyl-CoA transferase (FRC [EC 2.8.3.16]), oxalyl-CoA decarboxylase (OXC [EC 4.1.1.8]), and an oxalate:formate antiporter (OXLT). The genes *frc* and *oxc* are found as operons in many OxBs (Khammar et al. 2009). The second model is *via* aerobic metabolism, where oxalate is used as the energy source through carbohydrate metabolism or serine biosynthesis. The key players in this pathway are the enzymes glyoxylate carboligase (GCL [EC 4.1.1.47]) and hydroxypyruvate reductase [EC 1.1.1.29].

In this study, using a collection of bacteria (Bang et al. 2018) from the desert and drylands of the Middle East, we scouted for bacteria that possess the necessary toolkit of enzymes to degrade and assimilate oxalate (Additional Supplementary File 1). For this purpose, we identified 20 enzymes, including one transporter, and built a database of high-quality representative enzymes involved in all aspects of oxalate metabolism (Supplementary Table 1). We focused and validated oxalotrophy prediction on the family of Pseudomonads and revealed the genetic basis for the ability and inability of distinct strains to grow on oxalate.

## Methods

### Oxalate media growth test

All bacteria were tested on Schlegel mineral medium (Aragno and Schlegel 1992) containing calcium oxalate (Braissant, Verrecchia, and Aragno 2002). The first layer contains all components of Schlegel media: Na_2_HPO_4_×12H_2_O 9.0 g/l, KH_2_PO_4_ 1.5 g/l, NH_4_Cl 1.0 g/l, MgSO_4_×7H_2_O 0.2 g/l, ammoniacal ferric citrate 0.005 g/l, CaCl_2_ 0.01 g/l, ZnSO_4_×7 H_2_O 50 µg/l, MnCl_2_×4H_2_O 15 µg/l, boric acid 150 µg/l, CoCl_2_×6 H_2_O 100 µg/l, CuCl_2_×2H_2_O 50 µg/l, NiCl_2_×6H_2_O 10 µg/l, NaMoO_4_×2H_2_O 15 µg/l and 15 g/l agar. The second layer consists of Schlegel media containing 4g/l calcium oxalate monohydrate. The bacteria were grown at 28°C for two days before visual evaluation. As a control, the bacterial strains were grown at 28°C on LB agar media (Sigma-Aldrich).

### Identification of enzymes of oxalate metabolism

The enzymes associated with oxalate metabolism were identified in the literature and the BioCyc database (Karp et al. 2017). Twenty enzymes were involved in oxalate biosynthesis, assimilation, and degradation. We created a local protein sequence database by choosing the best-reviewed, preferably swiss-prot sequences (Consortium 2024) belonging to bacteria. When multiple sequences were available, all were included to capture sequence diversity. The accession for the sequences used for the database is shown in Supplementary Table 1.

The genomes for all the strains were sequenced (The HirtLab, KAUST) and assembled in-house with Spades v3.15.5 (Prjibelski et al. 2020). The assemblies were validated for completeness and contamination with checkM (Parks et al. 2015) and taxonomic assignment using ANI (Ciufo et al. 2018). The assemblies thus obtained were annotated using prokka v1.14.6 (Seemann 2014). The sequences less than 200bp were filtered out to eliminate the noise during annotation.

Next, protein sequences thus obtained from 536 strains were analyzed to construct the presence/absence and count number matrix for the identified enzymes. A ‘hit’ was defined by balancing both the sensitivity and specificity (Pearson 2013) of the BLAST hits (Altschul et al. 1990). Briefly, the hits were filtered with the parameters: E-value 1E-3, bitscore >= 40, percentage identity >= 30%, and at least 60% of query coverage. Phylogenetic relationships among the 536 bacterial strains were analyzed using the mashtree package v1.4 (Katz et al. 2019) to generate a phylogenetic tree, providing a broader context for the distribution of oxalate metabolism enzymes. The tree was visualized with iTOL v6 by mapping each species with their respective traits (Letunic and Bork 2024).

### Single nucleotide polymorphism (SNP) analysis

We performed SNP analysis using snippy v4.6.0 (Seemann 2015) to investigate the genetic variations between the genomes of interest. The variants were annotated with SnpEff (Cingolani et al. 2012) to classify high-impact and moderate-impact variations. The annotated sequences were further examined for the enzyme-related terms (EC numbers) to identify variations potentially affecting the enzyme function. Further, the selected protein sequences were aligned using Prank v.170427 (Löytynoja 2014) to assess sequence-level changes in terms of conservative and non-conservative amino acid substitutions.

## Results

### Strain Isolation and Collection

Our collection included 536 strains from 4 phyla and 81 genera isolated from 20 different desert host plants in the Middle East (Bang et al. 2018). Among the four phyla (Figure 1A), Pseudomonadota represents 57% of the isolates, with most of the genus belonging to *Pseudomonas*, while *Bacillota* occupies 20% (Figure 1B). Among the hosts, most strains were isolated from the *Avicennia marina* (mangrove biome), followed by the date palm tree *Phoenix dactylifera*. In the context of the study and the statistics, strains identified as “Pseudomonas” will be highlighted in the analyses.

**Figure 1:**
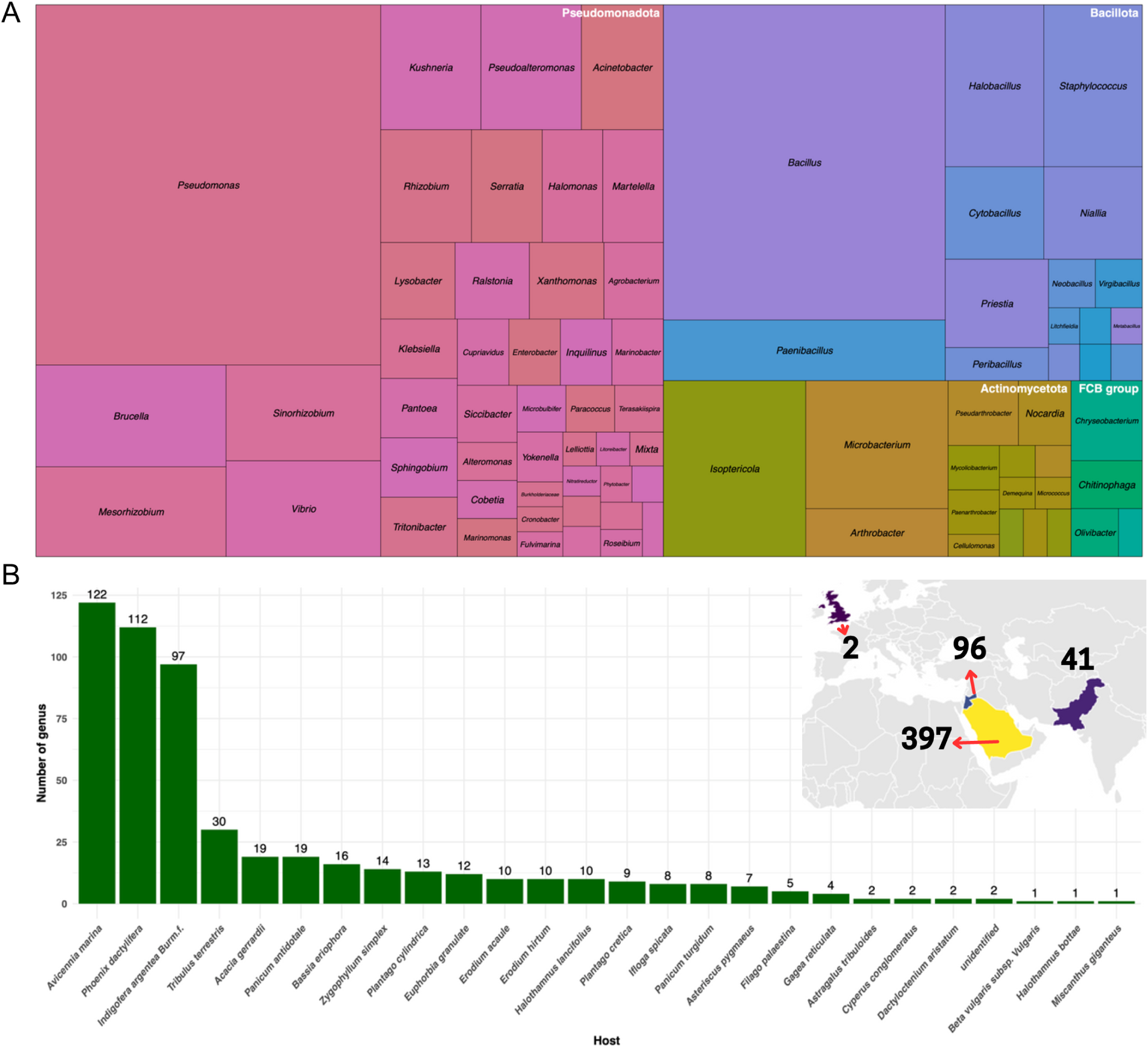
(A) Treemap showing the distribution of different genera grouped under Phylum. The box size represents the number of strains identified under each genus. (B) Distribution of genera under different hosts. The inset shows the country-wise distribution of isolated strains.

### Oxalate Metabolism

Plants and fungi are the main producers of oxalate. The biogenesis of oxalate is observed in very few bacteria, such as *P. fluorescens* ATCC 13525 (Hamel, Levasseur, and Appanna 1999) and *Burkholderia* species (Nakata and He 2010; Nakata 2011). Figure 2 shows the summarized reactions involved in oxalate metabolism. The bacteria can obtain oxalate *via* three routes, one as an external carbon source through the activity of an oxalate: formate antiporter under anaerobic metabolism. The second is as an intermediate of aerobic energy metabolism *via* the enzymatic activity of oxaloacetic hydrolase [EC 3.7.1.1], which catalyzes the conversion of oxaloacetate into oxalate and acetate. In route 3, the oxalate is formed by the breakdown of glyoxylate into oxalate and hydrogen peroxide by the enzymatic activity of glyoxylate oxidase.

**Figure 2:**
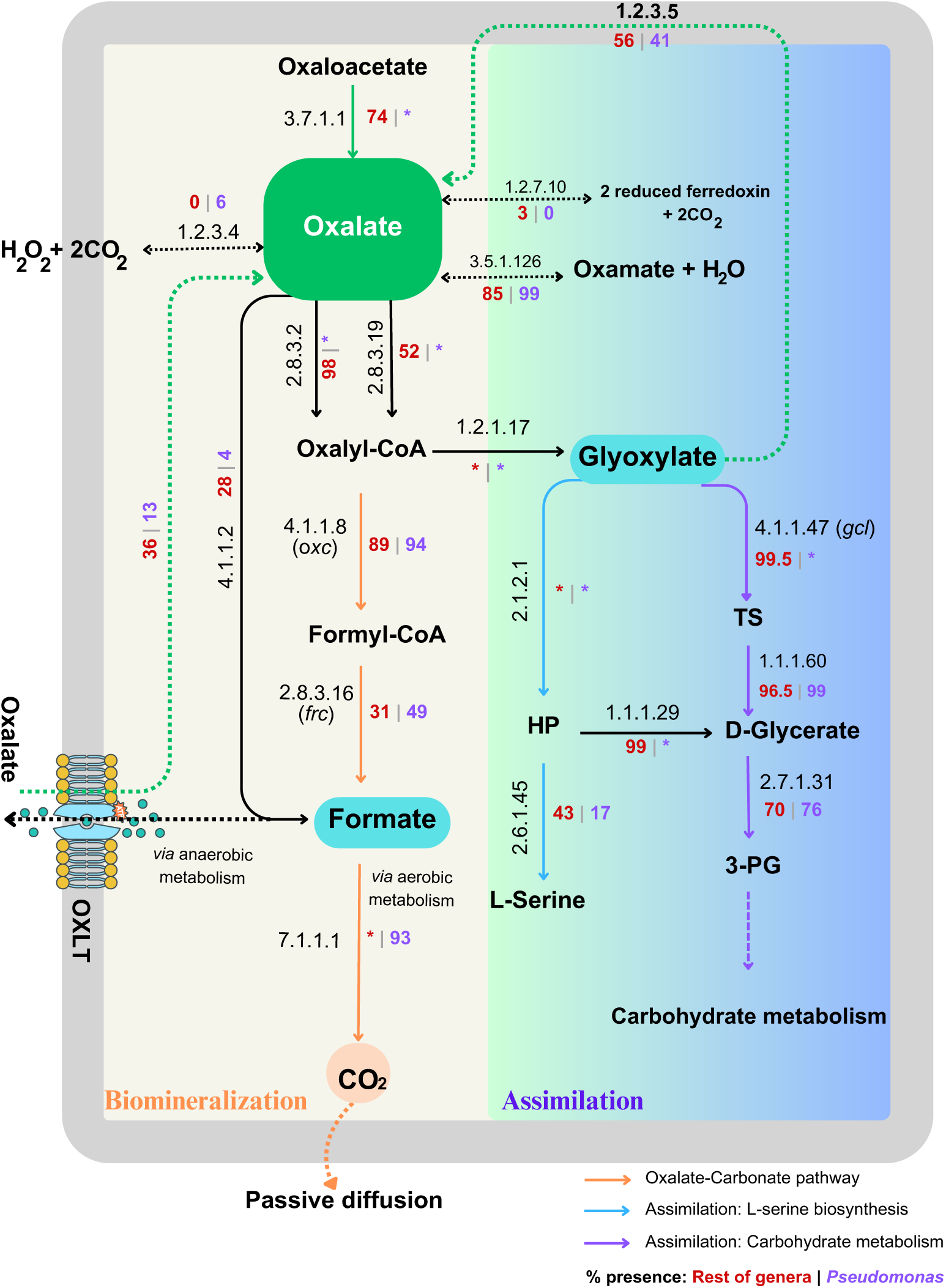
Reactions involved in oxalate metabolism. The pathways shown are in condensed form. i.e., only substrate and main intermediates are shown. Detailed reactions are described by Palmieri et al, 2019. The green arrow indicates the biosynthesis route. HP – hydroxy pyruvate, TS – Tartronate semialdehyde, PG-phosphoglycerate

The obtained oxalate enters either into the OCP or the assimilation pathway. In the OCP route, the oxalate is broken down to formate and carbon dioxide, contributing to inorganic carbon cycling. The carbon dioxide is excreted out of the cell by passive diffusion reaction, which is further used to precipitate minerals like calcium, thus leading to carbon sequestration. Briefly, the oxalate is converted to oxalyl-CoA by the enzymes oxalate CoA transferases (EC 2.8.3.2, EC 2.8.3.19]. The enzyme OXC (EC 4.1.1.8) catalyzes the formation of formyl-CoA using oxalyl-CoA as the substrate which is broken to formate by FRC (EC 2.8.3.16). Carbon dioxide is produced by the action of NAD-linked formate dehydrogenase (EC 7.1.1.1) on formate.

In the assimilation pathway, oxalate is utilized as the intermediate to synthesize amino acids or metabolize carbohydrates through the glycerate pathway. The intermediate Oxalyl-CoA is converted to glyoxylate catalyzed by Oxalyl-CoA reductase (EC 1.2.1.17). At this point, glyoxylate is the junction that determines the fate of the oxalate metabolism. Under the action of glyoxylate oxidase (EC 1.2.3.5), glyoxylate can be reversibly converted back to oxalate. Alternatively, the enzymes serine hydroxymethyltransferase (EC 2.1.2.1) and serine—glyoxylate transaminase (EC 2.6.1.45) lead to the biosynthesis of L-Serine.

Finally, to metabolize carbohydrates, glyoxylate is converted to 3-phosphoglycerate. In short, the enzyme glyoxylate carboligase (EC 4.1.1.47 – GCL) breaks the glyoxylate into tartronic semialdehyde, which is converted to D-glycerate by the action of a reductase (EC 1.1.1.60). In the last step, glycerate is converted into 3-PG by the action of glycerate 3-kinase (EC 2.7.1.31). Thus, 3-PG enters energy production through glycolysis and the Calvin cycle. These metabolic routes highlight the versatility of oxalate utilization in bacteria, linking it to both energy production and biosynthesis.

### Genus-wide distribution of enzymes

To analyze the genus-wide distribution of enzymes, we performed the presence-absence analysis of all the enzymes involved in the oxalate metabolism (Figure 3). The occurrence of each enzyme was calculated as a percentage for each sampled genus, providing an overview of their distribution. As depicted in the heatmap, the highly conserved enzymes are clustered together, whereas the less conserved enzymes are grouped towards the right side. The enzymes involved in the assimilation pathway (Figure 2) are highly conserved across all genera and participate in anaplerotic reactions, including examples such as 1.2.1.17, 2.8.3.2, 2.1.2.1, 1.1.1.29, GCL, and 1.1.1.60. Conversely, less conserved enzymes like 2.6.1.45, FRC, and 4.1.1.20 play crucial roles in the production of key metabolites such as L-serine, formate, and oxalate. These less conserved enzymes can, therefore, be characterized as trait specific. Additionally, clustering based on the presence or absence of the enzymes shows three clusters, ranging from ubiquitous presence to rare occurrence (Supplementary Figure 1).

**Figure 3:**
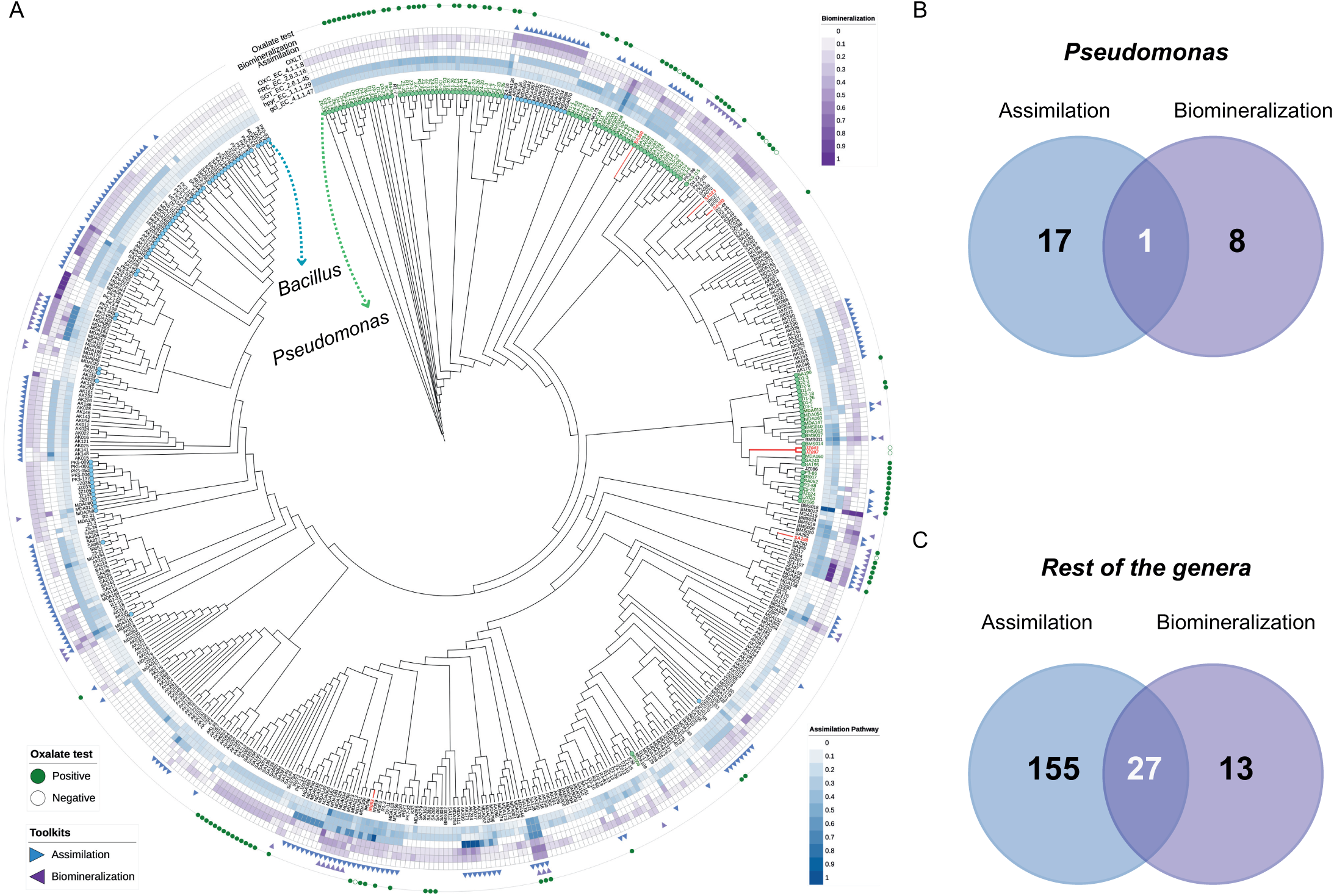
A) Copy numbers of enzymes in heatmap for assimilation and biomineralization toolkits. B) Venn diagram of strains in the Pseudomonas group that harbors the categorized enzyme repertoire. C) Venn diagram of strains in the rest of the genera. [Strain name: P3-86 *alias* JZ860]

The copy number distribution for enzymes analyzed is shown in Supplementary Figure 2. The tiles were arranged in descending order of copy number sums, spreading outwards of the circular tree. The first few enzymes near leaf labels show a consistent pattern across all the samples, which could indicate functional specialization or genomic similarity. On the other hand, the pattern is more sporadic towards the outer lanes, suggesting potential metabolic flexibility.

Among the genera studied, the FCB group was found to have several holes, even within the highly conserved category. However, this may be attributed to the small sample size of the FCB group, i.e., the total number of strains under the FCB group is eleven compared to 108 Pseudomonas and 78 Bacillus.

Overall, the percentage distribution of enzymes on the reaction pathways reveals clear preferential routes of oxalate metabolism among the genera. The Pseudomonas species prefers to assimilate the oxalate via carbohydrate metabolism rather than L-serine biosynthesis. 83% of the Pseudomonas group exhibit a propensity to produce D-glycerate either with tartronic semialdehyde or hydroxy pyruvate as an intermediate. On the other hand, the rest of the genera exhibit similar preferences for both carbohydrate and amino acid biosynthesis.

### Pathways of Oxalotrophy

We categorized the enzymes involved in oxalotrophy into pathway toolkits. The enzymes OXC (EC 4.1.1.8), FRC (EC 2.8.3.16), and the transporter OXLT were grouped under the biomineralization pathway. Similarly, the enzymes GCL (EC 4.1.1.47), hydroxy pyruvate reductase (EC 1.1.1.29), and serine-glyoxylate transaminase (EC 2.6.1.45) were classified as part of the assimilation pathway. We hypothesized that the completeness of these pathways would confer the ability of oxalotrophy to bacteria. To test our hypothesis, a genomic signature analysis was carried out to determine the presence of categorized pathways mentioned above. Figure 4A shows the copy number distribution of enzymes belonging to the categorized pathways.

**Figure 4:**
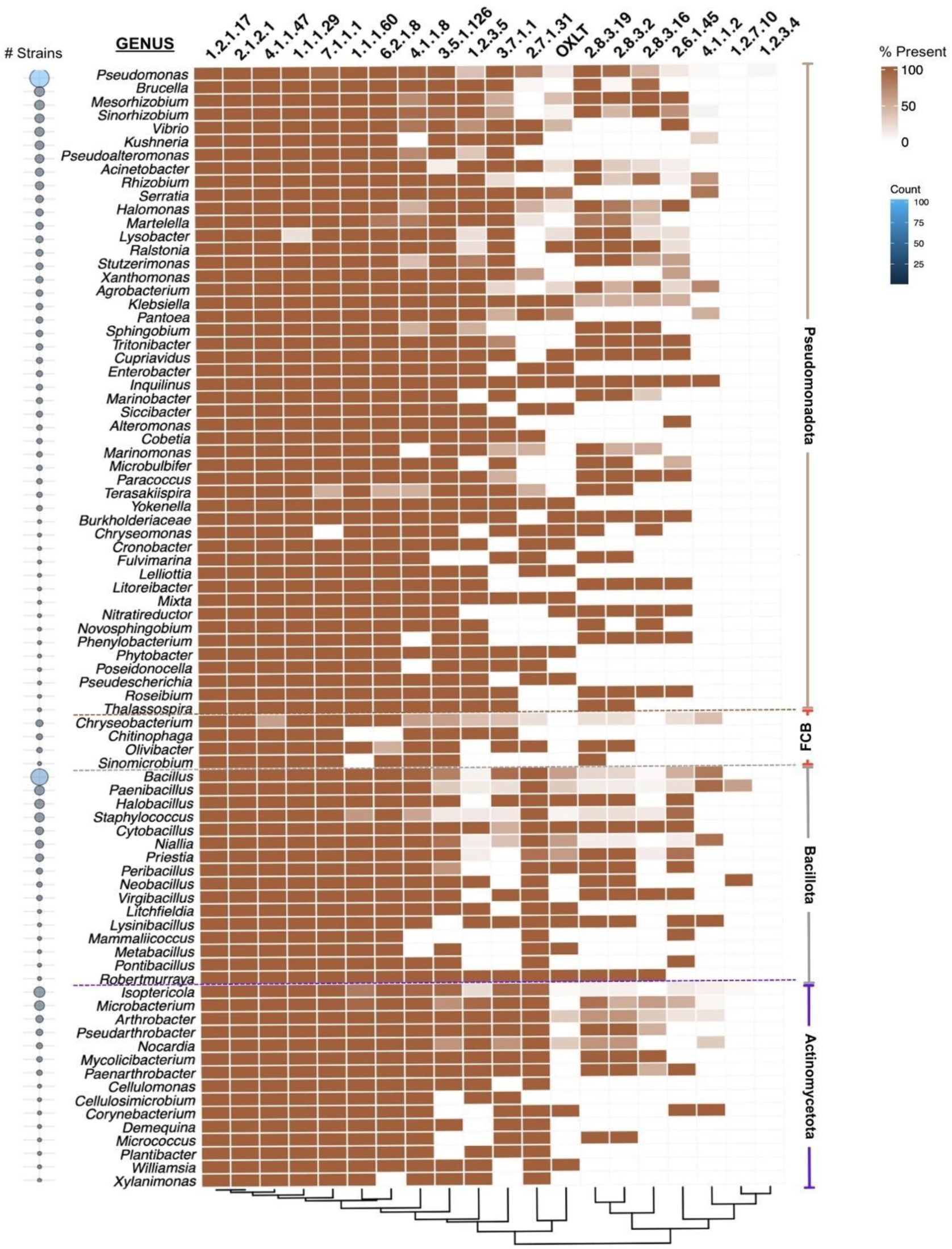
Heatmap showing the genus-wise percentage distribution of the enzymes involved in oxalate metabolism. For each phylum, the genus was sorted in descending order of number of strains identified (bubble in the left).

In our study, nine Pseudomonas strains (JZ006, JZ042, JZ091, JZ114, JZ134, MDA012, PK3-6, PK5-115, PK5-23) were identified with the capacity for biomineralization (Figure 4B). Interestingly, these strains lacked the enzyme EC 2.6.1.45, which is essential for the L-Serine biosynthesis. On the other hand, eighteen strains were identified with the complete enzyme set required for oxalate assimilation. Only one strain, *Pseudomonas sp.* MD012, isolated from the date palm *Phoenix dactylifera,* contained both the pathways to metabolize oxalate, showcasing its unique metabolic versatility.

In contrast, 40 and 182 strains of other genera contained enzyme sets to achieve biomineralization and assimilation, respectively (Figure 4C). In theory, 27 strains could convert oxalate into CO_2_ and assimilate it to L-Serine. Like in *Pseudomonas*, the strains that could perform only biomineralization lacked the key enzyme serine–glyoxylate transaminase EC 2.6.1.45. These findings underscore the critical role of enzyme EC 2.6.1.45 in oxalate assimilation. The strains capable of oxalate assimilation were associated with diverse hosts, including mangrove biomes. As a further validation, we tested several strains *in vitro* for their ability to grow on media supplemented with oxalate as the sole organic carbon source.

### Accumulation of SNPs in the key enzyme

Among the tested clades, one of the subclades belonging to *Pseudomonas* (Figure 5A) revealed an interesting pattern where all the tested strains grew on oxalate media, except for JZ097 and JZ043 (Figure 5B). To test whether mutations in one or more key enzymes within oxalate assimilation or biomineralization pathways might be the reason for the observed trait, we performed SNP analysis using *P. spp.* JZ024 (Eida et al. 2018; 2019), a species with a high-quality and complete genome as a reference within the subclade. The SNP analyses for all the tested genus are shown in Supplementary Table 2. Interestingly, the oxalate-negative strains were found to have missense SNPs in the enzyme glyoxylate carboligase (EC 4.1.1.47). This enzyme is involved in the conversion of glyoxylate to tartronate semialdehyde, which further proceeds to carbohydrate metabolism. The multiple-sequence alignment showed a clear difference in mosaic patterns between the groups (Figure 5A). In addition, more non-conservative substitutions that affect the protein function were observed compared to conserved substitutions.

**Figure 5:**
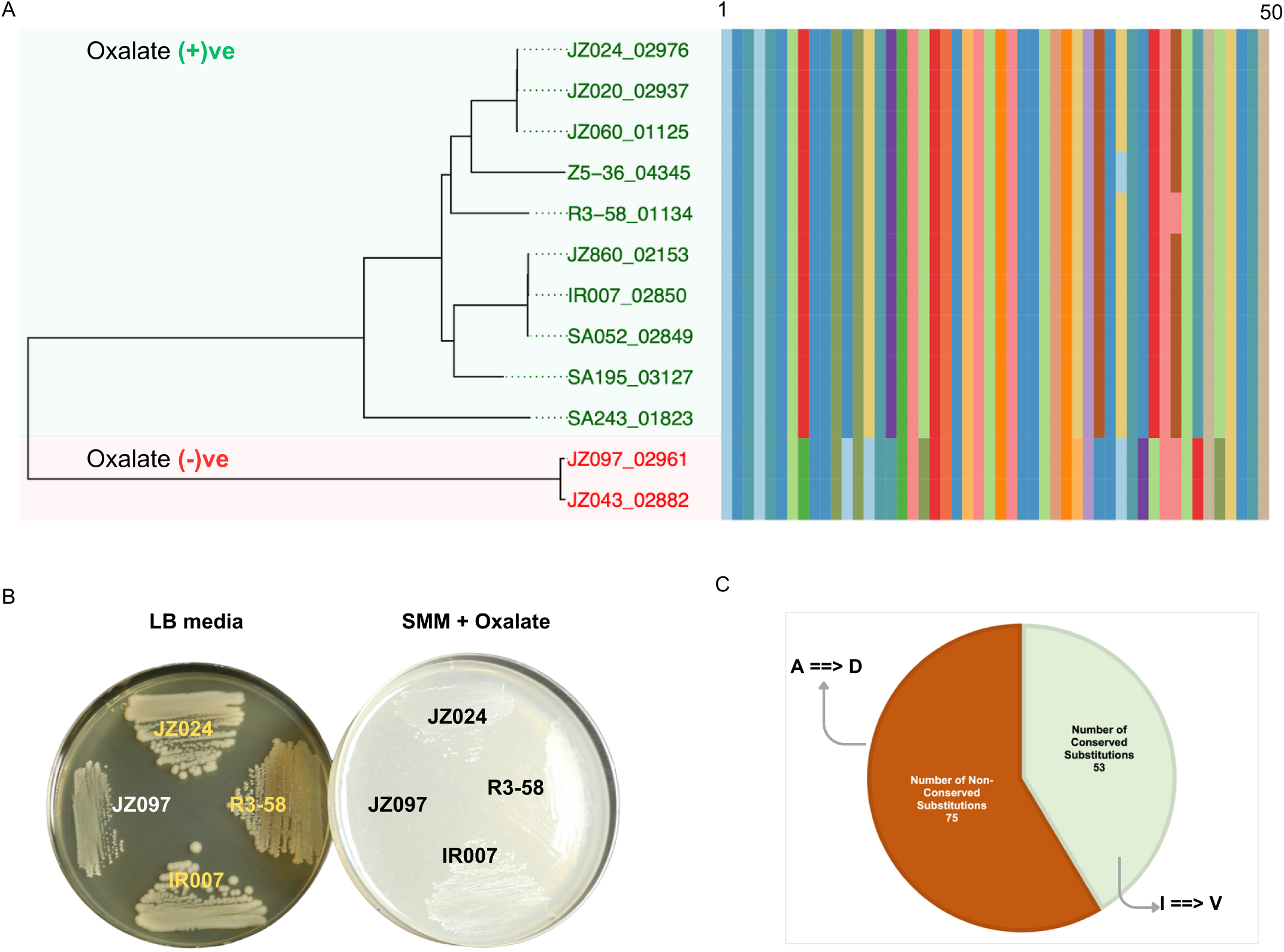
The Pseudomonas group sub-node tested for oxalate in the growth medium (A). The multiple sequence alignment between oxalate negative and positive strains. B) Strains tested on LB media and Schlegel mineral media supplemented with exogenous oxalate. C) Distribution of conserved and non-conserved substitutions.

A notable non-conservative and known detrimental substitution observed was Alanine to Aspartate (A to D). A study published in 1996 investigated the effect of substituting Alanine 128 with Aspartic Acid in the β-subunit of the F₀F₁-ATPase from *Escherichia coli*. This mutation abolished the dimerization of the β-subunit, which is crucial for the proper assembly and function of the ATPase complex (Howitt et al. 1996). Similarly, the most frequent conservative substitution observed was Isoleucine to Valine (I to V). Although the mutation is conservative, the position it appears can cause significant disruption in protein structure. In *E. coli* testing the effects of isoleucine-to-valine and valine-to-isoleucine mutations in thioredoxin resulted in changes in stability of the protein (Godoy-Ruiz et al. 2005). Thus, the mutations we observed in the GCL could result in impaired enzyme function, leading to the inability to grow in oxalate media.

## Discussion

The oxalate-carbonate pathway (OCP), facilitated by bacteria, represents a promising mechanism for long-term carbon sequestration in terrestrial ecosystems. This process plays a pivotal role in carbon cycling and contributes to soil carbon storage, offering a natural solution for mitigating atmospheric CO₂ levels. Central to bacterial oxalate metabolism are three key enzymes: formyl-CoA transferase, oxalyl-CoA decarboxylase, and the formate/oxalate reverse transporter (OxlT). These enzymes drive the conversion of oxalate into CO₂, which subsequently precipitates as calcium carbonate, thereby locking carbon in a stable mineral form (Daniel et al. 2021; Xia et al. 2024). In our study, we identified several bacterial isolates harboring these critical enzymes. Two enzymes, oxalate decarboxylase [EC 4.1.1.2] and oxalate oxidase [EC 1.2.3.4], are reported to oxidize oxalate primarily in fungi and lower eukaryotes (Cowan et al. 2024). Currently, available enzymatic methods for the quantitation of oxalate are based on hydrogen peroxide production via oxalate degradation using EC 1.2.3.4. We identified six Pseudomonas species, all isolated from Wadi rum, Jordan, to harbor these enzymes exclusively. Additionally, four Pseudomonas species (BMS010, BMS012, BMS014, and BMS017) from Al Ula, Saudi Arabia, were found to possess EC 4.1.1.2, suggesting distinct metabolic adaptations influenced by the environment or by horizontal gene transfer from other organisms such as fungi or lower eukaryotes. Incidentally, both these regions belong to the same geographical stretch and share properties of soil and ecology.

The inability of certain bacterial strains to grow on oxalate media, as observed in our results, highlights intriguing variability in the oxalotrophic potential among different bacteria. Our findings suggest that the accumulation of SNPs in key enzymes might be a reason for the inability of strains to utilize oxalate. The EC 4.1.1.47 protein sequences coding for glyoxylate carboligase of non-oxalotrophic strains JZ043 and JZ097 (Eida et al. 2019) were distinct from those of other strains, suggesting potential functional divergence related to oxalate metabolism. These findings underscore significant variability in the oxalotrophic potential among bacterial populations, indicating evolutionary and ecological factors influencing this trait.

As mentioned before, oxalic acid is a key root exudate, shaping rhizosphere microbial communities by recruiting beneficial oxalotrophic bacteria like *Burkholderia*, which promote plant growth and colonization. Oxalotrophy reduces oxalic acid levels, limiting fungal pathogenicity and providing plant protection (Kost et al. 2014). Studies show that oxalotrophic bacteria, such as *Pseudomonas spp*., reduce fungal infections in crops by 30–75% (Dickman and Chet 1998; Schoonbeek et al. 2007), highlighting their potential for sustainable agricultural applications. Furthermore, OxB can help mitigate some of the challenges faced in desert agriculture. For instance, their mineral weathering ability can release essential nutrients such as calcium and magnesium, enriching the nutrient pool in coarse-textured soil. OxB can also break down calcium carbonate into more bioavailable forms, improving root penetration and water infiltration. Further, OxB chelates toxic metals and ions in saline soils, which can reduce soil salinity and alkalinity stress on plants. Due to their broad environmental benefits, our work lays a crucial foundation for harnessing oxalotrophic bacteria in soil carbon sequestration strategies, especially in arid and semi-arid ecosystems.

## Conclusion

Our study highlights the metabolic versatility of oxalotrophic bacteria and their critical role in carbon cycling, soil health, and plant-microbe interactions. The identification of distinct metabolic pathways, along with genomic variability in key enzymes, underscores the adaptive nature of oxalotrophy. The ability of these bacteria to modulate oxalate levels has significant implications for soil carbon sequestration and sustainable agriculture, particularly in arid environments. Harnessing oxalotrophic bacteria could offer innovative strategies for enhancing soil resilience and mitigating climate change, warranting further research into their functional stability and ecological impact in natural ecosystems.

## Author contribution

H.H. conceived the idea presented. A.P. developed the concept, performed bioinformatics analyses, and wrote the manuscript. S.P. performed the initial study. K.S. contributed to the Introduction and Discussion sections. M.M.S. provided the strains presented in the manuscript and contributed to the conclusion. K.F., T.S., and D.R. performed experiments to test the strains. H.H., A.P., K.S., M.S., K.T., and T.S. proofread the final version of the manuscript.

## Acknowledgment

The authors thank all members of the Hirt lab, the CDA management team, and the Bioscience Core Labs at KAUST for their technical assistance and help with many aspects of this work.

---

**Supplementary Table 1.**
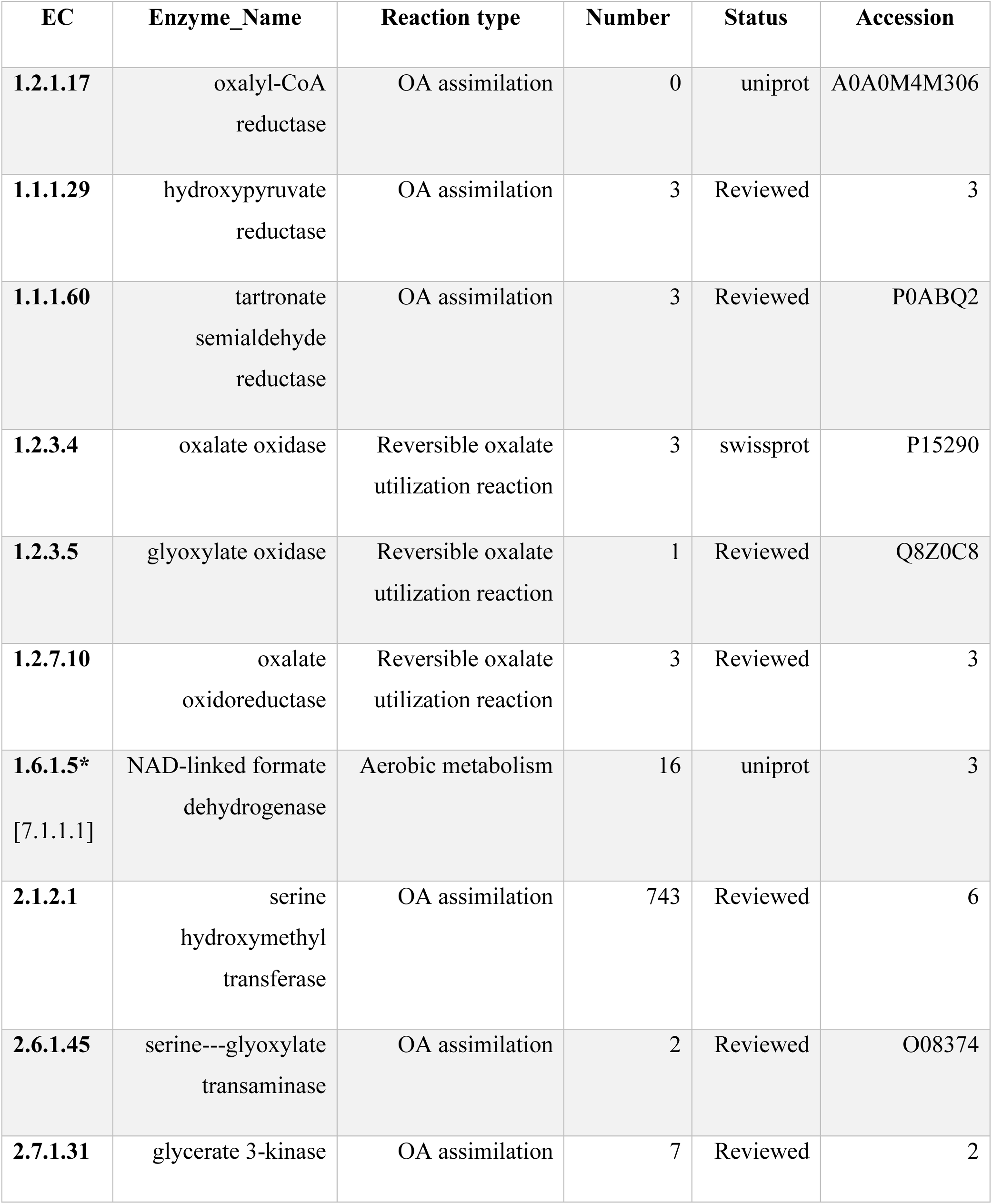
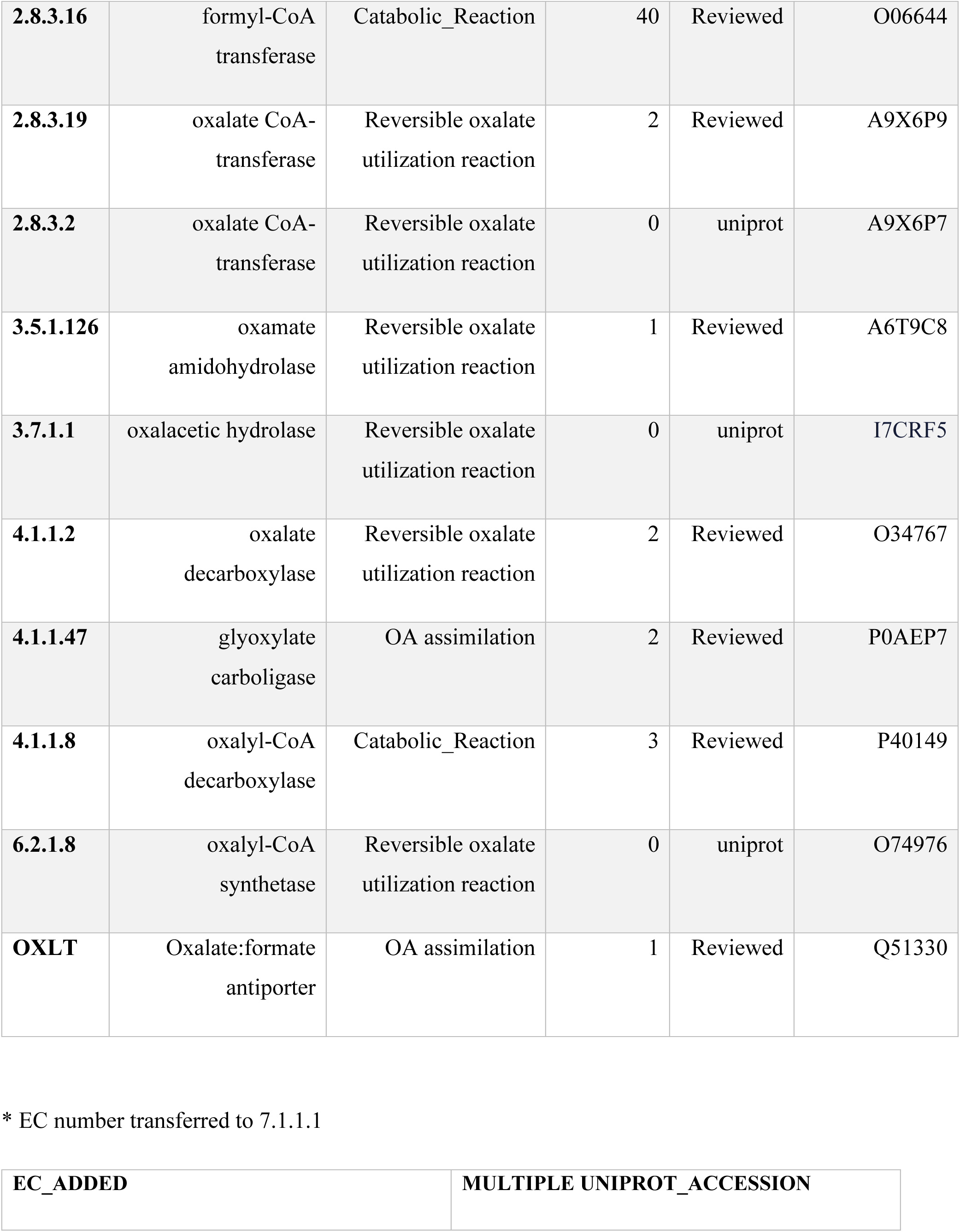
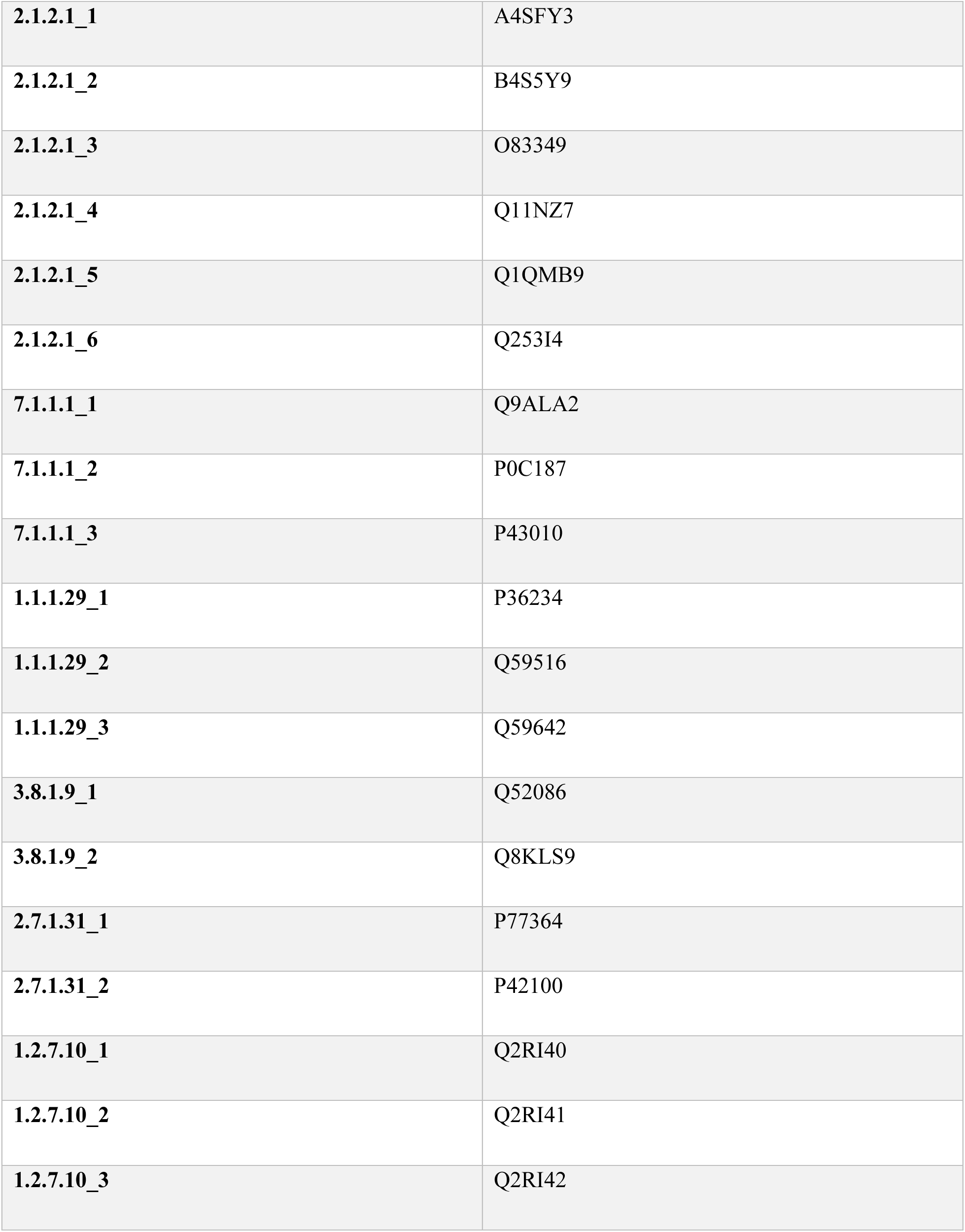

| **EC** | **Enzyme_Name** | **Reaction type** | **Number** | **Status** | **Accession** |
| --- | --- | --- | --- | --- | --- |
| **1.2.1.17** | oxalyl-CoA<br>reductase | OA assimilation | 0 | uniprot | A0A0M4M306 |
| **1.1.1.29** | hydroxypyruvate<br>reductase | OA assimilation | 3 | Reviewed | 3 |
| **1.1.1.60** | tartronate<br>semialdehyde<br>reductase | OA assimilation | 3 | Reviewed | P0ABQ2 |
| **1.2.3.4** | oxalate oxidase | Reversible oxalate<br>utilization reaction | 3 | swissprot | P15290 |
| **1.2.3.5** | glyoxylate oxidase | Reversible oxalate<br>utilization reaction | 1 | Reviewed | Q8Z0C8 |
| **1.2.7.10** | oxalate<br>oxidoreductase | Reversible oxalate<br>utilization reaction | 3 | Reviewed | 3 |
| **1.6.1.5\***<br>[7.1.1.1] | NAD-linked formate<br>dehydrogenase | Aerobic metabolism | 16 | uniprot | 3 |
| **2.1.2.1** | serine<br>hydroxymethyl<br>transferase | OA assimilation | 743 | Reviewed | 6 |
| **2.6.1.45** | serine---glyoxylate<br>transaminase | OA assimilation | 2 | Reviewed | O08374 |
| **2.7.1.31** | glycerate 3-kinase | OA assimilation | 7 | Reviewed | 2 |
| **2.8.3.16** | formyl-CoA<br>transferase | Catabolic_Reaction | 40 | Reviewed | O06644 |
| **2.8.3.19** | oxalate CoA-transferase | Reversible oxalate<br>utilization reaction | 2 | Reviewed | A9X6P9 |
| **2.8.3.2** | oxalate CoA-transferase | Reversible oxalate<br>utilization reaction | 0 | uniprot | A9X6P7 |
| **3.5.1.126** | oxamate<br>amidohydrolase | Reversible oxalate<br>utilization reaction | 1 | Reviewed | A6T9C8 |
| **3.7.1.1** | oxalacetic hydrolase | Reversible oxalate<br>utilization reaction | 0 | uniprot | I7CRF5 |
| **4.1.1.2** | oxalate<br>decarboxylase | Reversible oxalate<br>utilization reaction | 2 | Reviewed | O34767 |
| **4.1.1.47** | glyoxylate<br>carboligase | OA assimilation | 2 | Reviewed | P0AEP7 |
| **4.1.1.8** | oxalyl-CoA<br>decarboxylase | Catabolic_Reaction | 3 | Reviewed | P40149 |
| **6.2.1.8** | oxalyl-CoA<br>synthetase | Reversible oxalate<br>utilization reaction | 0 | uniprot | O74976 |
| **OXLT** | Oxalate:formate<br>antiporter | OA assimilation | 1 | Reviewed | Q51330 |

\* EC number transferred to 7.1.1.1

|  |  |
| --- | --- |
| **EC_ADDED** | **MULTIPLE UNIPROT_ACCESSION** |
| **2.1.2.1_1** | A4SFY3 |
| **2.1.2.1_2** | B4S5Y9 |
| **2.1.2.1_3** | O83349 |
| **2.1.2.1_4** | Q11NZ7 |
| **2.1.2.1_5** | Q1QMB9 |
| **2.1.2.1_6** | Q253I4 |
| **7.1.1.1_1** | Q9ALA2 |
| **7.1.1.1_2** | P0C187 |
| **7.1.1.1_3** | P43010 |
| **1.1.1.29_1** | P36234 |
| **1.1.1.29_2** | Q59516 |
| **1.1.1.29_3** | Q59642 |
| **3.8.1.9_1** | Q52086 |
| **3.8.1.9_2** | Q8KLS9 |
| **2.7.1.31_1** | P77364 |
| **2.7.1.31_2** | P42100 |
| **1.2.7.10_1** | Q2RI40 |
| **1.2.7.10_2** | Q2RI41 |
| **1.2.7.10_3** | Q2RI42 |

**Supplementary Table 2.**

| **Genus** | **Strains tested** | **Mutation in Key enzymes** |
| --- | --- | --- |
| *Pseudomonas* | Shown in Figure 5 | EC 4.1.1.47 |
| *Sinorhizobium* | IR002, **IR052**,IR059, **IR066** * | EC 2.6.1.45, EC 4.1.1.47 |
| *Ralstonia* | SA290, **SA292** *, **SA288**, SA306 | - |
| *Serratia* | IR057, **SA023**, **SA093**,IR027,SA070, **SA088** * | - |

ST2: **Red** – Negative strains; \*- Used as genome reference for SNP analysis.

**Supplementary Figure 1.**
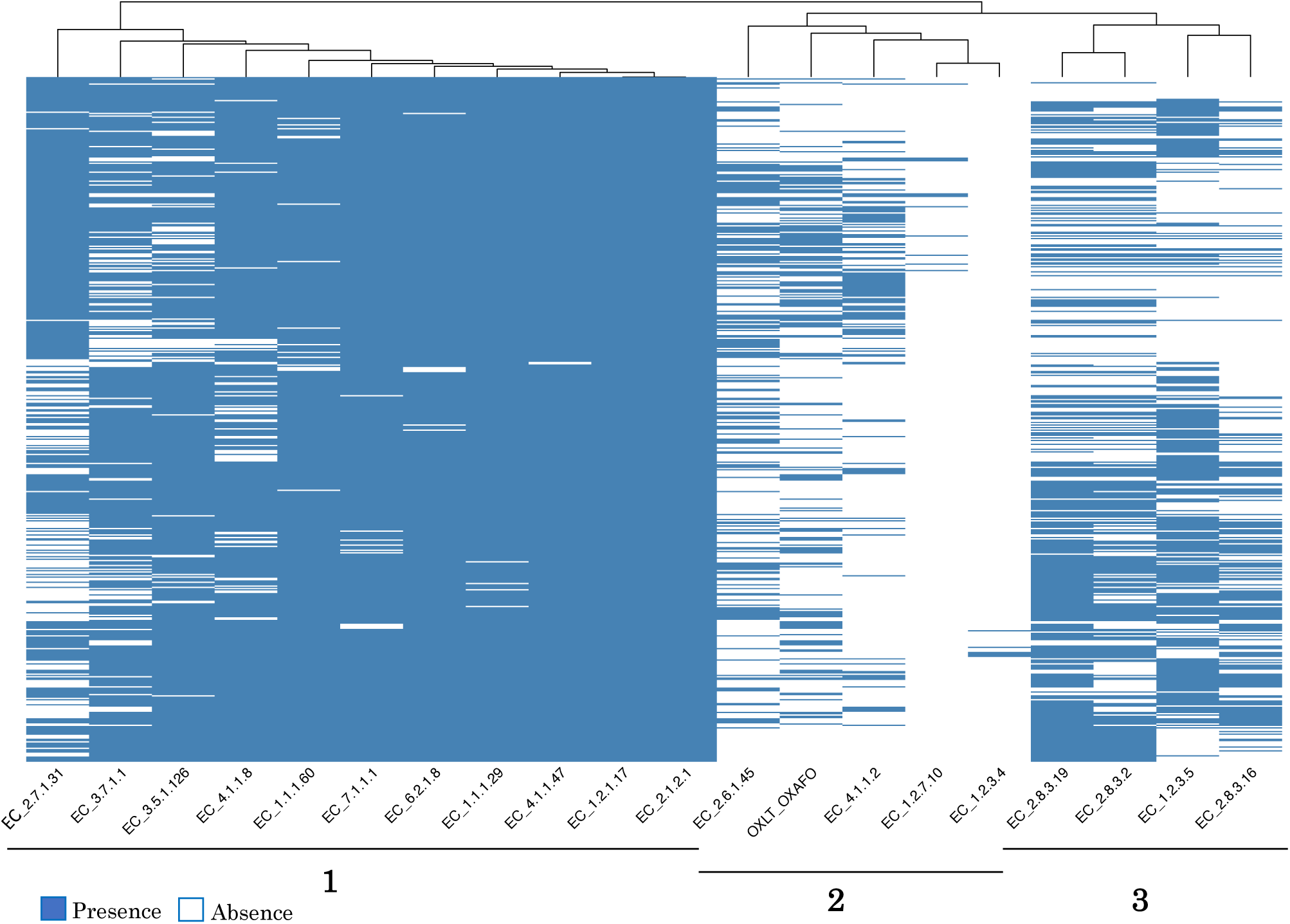
SF1: Heatmap of presence or absence of Enzymes across all the strains.

**Supplementary Figure 2:**
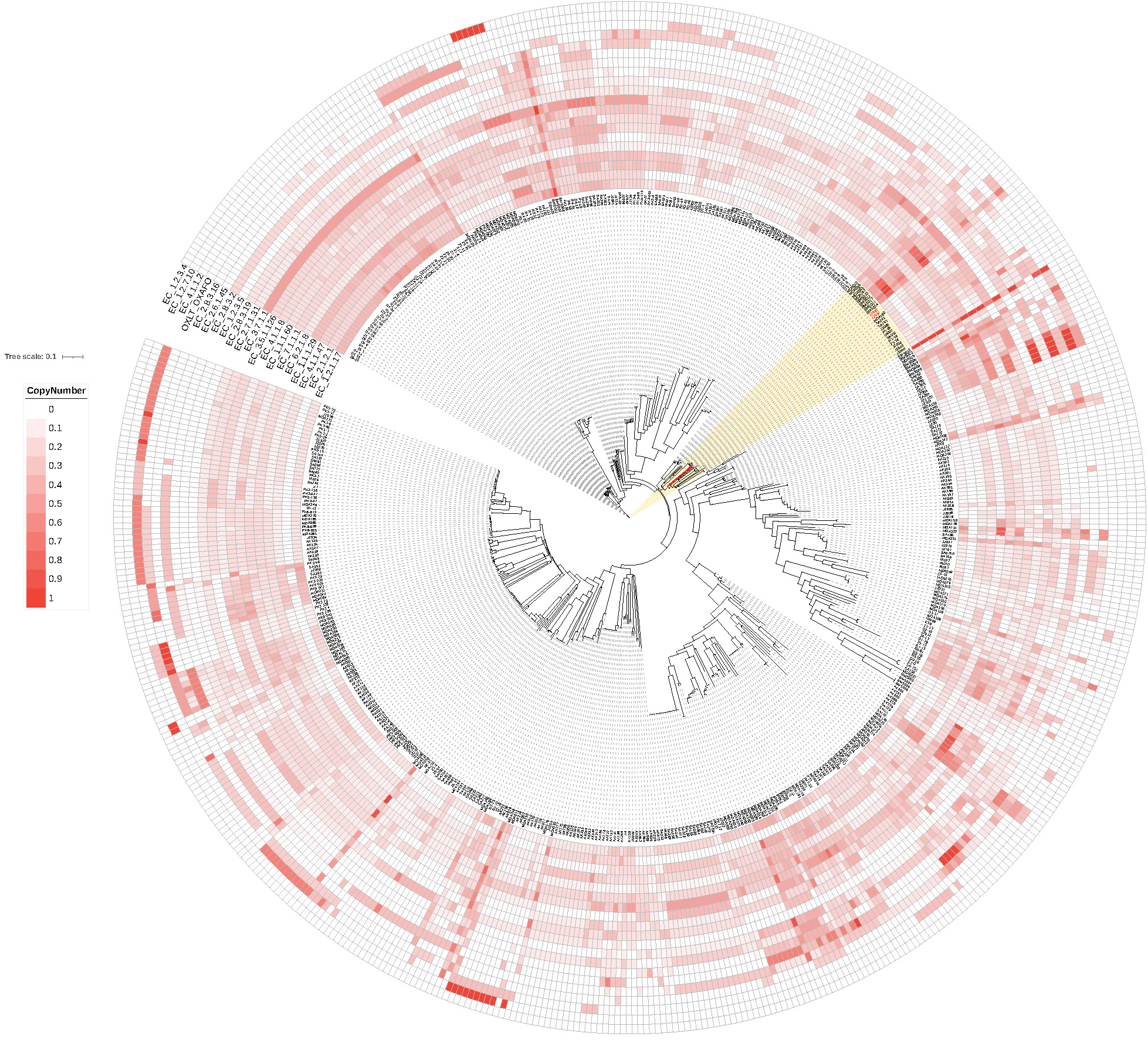
SF2: Copy number distribution of key enzymes involved in oxalate metabolism. Copy numbers are normalized for each node for visual representation. The highlighted clade shows the *Pseudomonas* subclade where strains tested negative (red) in oxalate media appear.

## References

Allison, Milton J., Karl A. Dawson, William R. Mayberry, and John G. Foss. 1985. “Oxalobacter Formigenes Gen. Nov., Sp. Nov.: Oxalate-Degrading Anaerobes That Inhabit the Gastrointestinal Tract.” Archives of Microbiology 141 (1): 1–7. 10.1007/BF00446731.

Altschul, Stephen F., Warren Gish, Webb Miller, Eugene W. Myers, and David J. Lipman. 1990. “Basic Local Alignment Search Tool.” Journal of Molecular Biology 215 (3): 403–10. 10.1016/S0022-2836(05)80360-2.

Aragno, M., and H. G. Schlegel. 1992. “The Mesophilic Hydrogen-Oxidizing (Knallgas) Bacteria.” 344–84. New York: Springer-Verlag Inc.

Aragno, M., and É Verrecchia. 2012. “The Oxalate-Carbonate Pathway: A Reliable Sink for Atmospheric CO2 Through Calcium Carbonate Biomineralization in Ferralitic Tropical Soils.” 10.1007/978-94-007-2229-3_9.

Bang, Corinna, Tal Dagan, Peter Deines, Nicole Dubilier, Wolfgang J. Duschl, Sebastian Fraune, Ute Hentschel, et al. 2018. “Metaorganisms in Extreme Environments: Do Microbes Play a Role in Organismal Adaptation?” Zoology 127:1–19. 10.1016/j.zool.2018.02.004.

Braissant, O., É Verrecchia, and M. Aragno. 2002. “Is the Contribution of Bacteria to Terrestrial Carbon Budget Greatly Underestimated?” Naturwissenschaften 89:366–70. 10.1007/s00114-002-0340-0.

Cingolani, Pablo, Adrian Platts, Le Lily Wang, Melissa Coon, Tung Nguyen, Luan Wang, Susan J. Land, Xiangyi Lu, and Douglas M. Ruden. 2012. “A Program for Annotating and Predicting the Effects of Single Nucleotide Polymorphisms, SnpEff: SNPs in the Genome of Drosophila Melanogaster Strain W1118; Iso-2; Iso-3.” Fly 6 (2): 80–92. 10.4161/fly.19695.

Ciufo, Stacy, Sivakumar Kannan, Shobha Sharma, Azat Badretdin, Karen Clark, Seán Turner, Slava Brover, Conrad L. Schoch, Avi Kimchi, and Michael DiCuccio. 2018. “Using Average Nucleotide Identity to Improve Taxonomic Assignments in Prokaryotic Genomes at the NCBI.” International Journal of Systematic and Evolutionary Microbiology 68 (7): 2386–92. 10.1099/ijsem.0.002809.

Consortium, The UniProt. 2024. “UniProt: The Universal Protein Knowledgebase in 2025.” Nucleic Acids Research 53 (D1): D609–17. 10.1093/nar/gkae1010.

Cowan, Don A, Darya Babenko, Ryan Bird, Alf Botha, Daniel O Breecker, Cathy E Clarke, Michele L Francis, et al. 2024. “Oxalate and Oxalotrophy: An Environmental Perspective.” Sustainable Microbiology 1 (1): qvad004. 10.1093/sumbio/qvad004.

Daniel, Steven L., Luke Moradi, Henry Paiste, Kyle D. Wood, Dean G. Assimos, Ross P. Holmes, Lama Nazzal, Marguerite Hatch, and John Knight. 2021. “Forty Years of Oxalobacter Formigenes, a Gutsy Oxalate-Degrading Specialist.” Applied and Environmental Microbiology 87 (18): e00544–21. 10.1128/AEM.00544-21.

Dickman, Martin B., and Iian Chet. 1998. “Biodegradation of Oxalic Acid: A Potential New Approach to Biological Control.” Soil Biology and Biochemistry 30 (8): 1195–97. 10.1016/S0038-0717(98)00018-2.

Duncan, Sylvia H., Anthony J. Richardson, Poonam Kaul, Ross P. Holmes, Milton J. Allison, and Colin S. Stewart. 2002. “*Oxalobacter Formigenes* and Its Potential Role in Human Health.” Applied and Environmental Microbiology 68 (8): 3841–47. 10.1128/AEM.68.8.3841-3847.2002.

Eida, Abdul Aziz, Hanin S. Alzubaidy, Axel de Zélicourt, Lukáš Synek, Wiam Alsharif, Feras F. Lafi, Heribert Hirt, and Maged M. Saad. 2019. “Phylogenetically Diverse Endophytic Bacteria from Desert Plants Induce Transcriptional Changes of Tissue-Specific Ion Transporters and Salinity Stress in Arabidopsis Thaliana.” Plant Science 280:228–40. 10.1016/j.plantsci.2018.12.002.

Eida, Abdul Aziz, Maren Ziegler, Feras F. Lafi, Craig T. Michell, Christian R. Voolstra, Heribert Hirt, and Maged M. Saad. 2018. “Desert Plant Bacteria Reveal Host Influence and Beneficial Plant Growth Properties.” PLOS ONE 13 (12): 1–20. 10.1371/journal.pone.0208223.

Ermer, Theresa, Lama Nazzal, Maria Clarissa Tio, Sushrut Waikar, Peter S. Aronson, and Felix Knauf. 2023. “Oxalate Homeostasis.” Nature Reviews Nephrology 19 (2): 123–38. 10.1038/s41581-022-00643-3.

Godoy-Ruiz, Raquel, Raul Perez-Jimenez, Beatriz Ibarra-Molero, and Jose M. Sanchez-Ruiz. 2005. “A Stability Pattern of Protein Hydrophobic Mutations That Reflects Evolutionary Structural Optimization.” Biophysical Journal 89 (5): 3320–31. 10.1529/biophysj.105.067025.

Hamel, Robert, Rémi Levasseur, and Vasu D. Appanna. 1999. “Oxalic Acid Production and Aluminum Tolerance in Pseudomonas Fluorescens.” Journal of Inorganic Biochemistry 76 (2): 99–104. 10.1016/S0162-0134(99)00120-8.

Hirt, Heribert, Hassan Boukcim, Marc Ducousso, and Maged M. Saad. 2023. “Engineering Carbon Sequestration on Arid Lands.” Trends in Plant Science 28 (11): 1218–21. 10.1016/j.tplants.2023.08.009.

Howitt, S. M., A. J. Rodgers, P. D. Jeffrey, and G. B. Cox. 1996. “A Mutation in Which Alanine 128 Is Replaced by Aspartic Acid Abolishes Dimerization of the B-Subunit of the F0F1-ATPase from Escherichia Coli.” The Journal of Biological Chemistry 271 (12): 7038–42. 10.1074/jbc.271.12.7038.

Karamad, Dina, Kianoush Khosravi-Darani, Amin Mousavi Khaneghah, and Aaron W. Miller. 2022. “Probiotic Oxalate-Degrading Bacteria: New Insight of Environmental Variables and Expression of the Oxc and Frc Genes on Oxalate Degradation Activity.” Foods 11 (18): 2876. 10.3390/foods11182876.

Karp, Peter D, Richard Billington, Ron Caspi, Carol A Fulcher, Mario Latendresse, Anamika Kothari, Ingrid M Keseler, et al. 2017. “The BioCyc Collection of Microbial Genomes and Metabolic Pathways.” Briefings in Bioinformatics 20 (4): 1085–93. 10.1093/bib/bbx085.

Katz, Lee S., Taylor Griswold, Shatavia S. Morrison, Jason A. Caravas, Shaokang Zhang, Henk C. den Bakker, Xiangyu Deng, and Heather A. Carleton. 2019. “Mashtree: A Rapid Comparison of Whole Genome Sequence Files.” Journal of Open Source Software 4 (44): 1762. 10.21105/joss.01762.

Khammar, N., Gaëtan Martin, K. Ferro, D. Job, M. Aragno, and É Verrecchia. 2009. “Use of the Frc Gene as a Molecular Marker to Characterize Oxalate-Oxidizing Bacterial Abundance and Diversity Structure in Soil.” Journal of Microbiological Methods 76 2:120–27. 10.1016/j.mimet.2008.09.020.

Kost, Thomas D., N. Stopnisek, Kirsty Agnoli, L. Eberl, and L. Weisskopf. 2014. “Oxalotrophy, a Widespread Trait of Plant-Associated Burkholderia Species, Is Involved in Successful Root Colonization of Lupin and Maize by Burkholderia Phytofirmans.” Frontiers in Microbiology 4:null. 10.3389/fmicb.2013.00421.

Kumar, Vinay, Mohammad Irfan, and Asis Datta. 2019. “Manipulation of Oxalate Metabolism in Plants for Improving Food Quality and Productivity.” Phytochemistry 158 (February):103–9. 10.1016/j.phytochem.2018.10.029.

Letunic, Ivica, and Peer Bork. 2024. “Interactive Tree of Life (iTOL) v6: Recent Updates to the Phylogenetic Tree Display and Annotation Tool.” Nucleic Acids Research 52 (W1): W78–82. 10.1093/nar/gkae268.

Löytynoja, Ari. 2014. “Phylogeny-Aware Alignment with PRANK.” *Methods in Molecular Biology (Clifton*, N.J.) 1079:155–70. 10.1007/978-1-62703-646-7_10.

Lux, Alexander, Jana Kohanová, and Philip J White. 2021. “The Secrets of Calcicole Species Revealed.” Journal of Experimental Botany 72 (4): 968–70. 10.1093/jxb/eraa555.

Nakata, Paul A. 2003. “Advances in Our Understanding of Calcium Oxalate Crystal Formation and Function in Plants.” Plant Science 164 (6): 901–9. 10.1016/S0168-9452(03)00120-1.

Nakata, Paul A. 2011. “The Oxalic Acid Biosynthetic Activity of Burkholderia Mallei Is Encoded by a Single Locus.” Microbiological Research 166 (7): 531–38. 10.1016/j.micres.2010.11.002.

Nakata, Paul A., and Cixin He. 2010. “Oxalic Acid Biosynthesis Is Encoded by an Operon in Burkholderia Glumae.” FEMS Microbiology Letters 304 (2): 177–82. 10.1111/j.1574-6968.2010.01895.x.

Naorem, Anandkumar, Somasundaram Jayaraman, Yash P. Dang, Ram C. Dalal, Nishant K. Sinha, Ch. Srinivasa Rao, and Ashok K. Patra. 2023. “Soil Constraints in an Arid Environment—Challenges, Prospects, and Implications.” Agronomy 13 (1). 10.3390/agronomy13010220.

Palmieri, Fabio, Aislinn Estoppey, Geoffrey L. House, Andrea Lohberger, S. Bindschedler, P. Chain, and P. Junier. 2019. “Oxalic Acid, a Molecule at the Crossroads of Bacterial-Fungal Interactions.” Advances in Applied Microbiology 106:49–77. 10.1016/bs.aambs.2018.10.001.

Parks, Donovan H., Michael Imelfort, Connor T. Skennerton, Philip Hugenholtz, and Gene W. Tyson. 2015. “CheckM: Assessing the Quality of Microbial Genomes Recovered from Isolates, Single Cells, and Metagenomes.” Genome Research 25 (7): 1043–55. 10.1101/gr.186072.114.

Pearson, William R. 2013. “An Introduction to Sequence Similarity (‘Homology’) Searching.” Current Protocols in Bioinformatics 42 (1): 3.1.1-3.1.8. 10.1002/0471250953.bi0301s42.

Pons, Sophie, S. Bindschedler, D. Sebag, P. Junier, É Verrecchia, and G. Cailleau. 2018. “Biocontrolled Soil Nutrient Distribution under the Influence of an Oxalogenic-Oxalotrophic Ecosystem.” Plant and Soil 425:145–60. 10.1007/s11104-018-3573-1.

Prjibelski, Andrey, Dmitry Antipov, Dmitry Meleshko, Alla Lapidus, and Anton Korobeynikov. 2020. “Using SPAdes De Novo Assembler.” Current Protocols in Bioinformatics 70 (1): e102. 10.1002/cpbi.102.

Schmalenberger, Achim, Achim Schmalenberger, Adele L. Duran, A. Bray, Jonathan Bridge, S. Bonneville, S. Bonneville, et al. 2015. “Oxalate Secretion by Ectomycorrhizal Paxillus Involutus Is Mineral-Specific and Controls Calcium Weathering from Minerals.” Scientific Reports 5:null. 10.1038/srep12187.

Schoonbeek, Henk-jan, Anne-Claude Jacquat-Bovet, Fabio Mascher, and Jean-Pierre Métraux. 2007. “Oxalate-Degrading Bacteria Can Protect Arabidopsis Thaliana and Crop Plants Against Botrytis Cinerea.” Molecular Plant-Microbe Interactions® 20 (12): 1535–44. 10.1094/MPMI-20-12-1535.

Seemann, Torsten. 2014. “Prokka: Rapid Prokaryotic Genome Annotation.” Bioinformatics 30 (14): 2068–69. 10.1093/bioinformatics/btu153.

Seemann, Torsten. 2015. “Snippy: Fast Bacterial Variant Calling from NGS Reads.” https://github.com/tseemann/snippy.

Xia, Dening, Wenjun Nie, Xiaofang Li, Roger D. Finlay, and Bin Lian. 2024. “Secondary Products and Molecular Mechanism of Calcium Oxalate Degradation by the Strain Azospirillum Sp. OX-1.” Scientific Reports 14:null. 10.1038/s41598-024-74939-8.

